# Hydrozoan sperm-specific H2B histone variants stabilize chromatin and block transcription without enhancing chromatin condensation

**DOI:** 10.1101/2021.08.30.458175

**Authors:** Anna Török, Martin JG Browne, Jordina C Vilar, Indu Patwal, Timothy Q DuBuc, Febrimarsa, Erwan Atcheson, Andrew Flaus, Uri Frank, Sebastian G Gornik

## Abstract

Many animals achieve sperm chromatin compaction and stabilisation during spermatogenesis by replacing canonical histones with sperm nuclear basic proteins (SNBPs) such as protamines. A number of animals including hydrozoan cnidarians and echinoid sea urchins lack protamines and have instead evolved a distinctive family of sperm-specific histone H2Bs (spH2Bs) with extended N-termini rich in SPKK-related motifs. Sperm packaging in echinoids such as sea urchins is regulated by spH2Bs and their sperm is negatively buoyant for fertilization on the sea floor. Hydroid cnidarians also package sperm with spH2Bs but undertake broadcast spawning and their sperm properties are poorly characterised. We show that sperm chromatin from the hydroid *Hydractinia* possesses higher stability than its somatic equivalent, with reduced accessibility of sperm chromatin to transposase Tn5 integration *in vivo* and to endonucleases *in vitro*. However, nuclear dimensions are only moderately reduced in mature *Hydractinia* sperm compared to other cell types. Ectopic expression of spH2B in the background of H2B knockdown resulted in downregulation of global transcription and cell cycle arrest in embryos without altering their nuclear density. Taken together, spH2B variants containing SPKK-related motifs act to stabilise chromatin and silence transcription in *Hydractinia* sperm without significant chromatin compaction. This is consistent with a contribution of spH2B to sperm buoyancy as a reproductive adaptation.

## Introduction

Male haploid sperm are produced from diploid progenitors. In this process, progenitor chromatin undergoes major remodelling to produce mature sperm chromatin exhibiting a high degree of condensation and stability compared with somatic chromatin (Sassone-Corsi, 2002; Ward and Coffey, 1991). This is achieved, in part, by sperm nuclear basic proteins (SNBPs) that replace somatic histones. SNBPs are structurally heterogeneous across species and are classified as histone-related, protamines, or protamine-related proteins (Török and Gornik, 2018).

Sequenced hydrozoan cnidarian genomes lack genes for protamines and instead encode a number of sperm-specific histone H2B variants (spH2Bs) that are structurally related to a protein family previously characterised in echinoid sea urchins (Ausio, 1999; Ausio et al., 1997; Busslinger and Barberis, 1985; Marzluff et al., 2006; Pérez-Montero et al., 2016; Poccia and Green, 1992; Rocchini et al., 1996; Török et al., 2016) (Fig. 1A). These histones include an extended N-terminal tail with up to 7 repeats of a SP[K/R][K/R] motif known to bind with high affinity to DNA minor grooves, preferentially at A/T rich sequences (Churchill and Suzuki, 1989; Khadake and Rao, 1997; Suzuki, 1989; Suzuki et al., 1993; Suzuki et al., 1990). Phosphorylation of the serine on the SPKK-related motifs in sea urchins inhibits DNA-binding (Green and Poccia, 1985; Poccia and Green, 1992). SPKK-related motif containing spH2Bs change the condensation status of sperm chromatin in echinoids and make chromatin inaccessible for transcription (Poccia and Green, 1992; Poccia et al., 1987).

**Figure 1:**
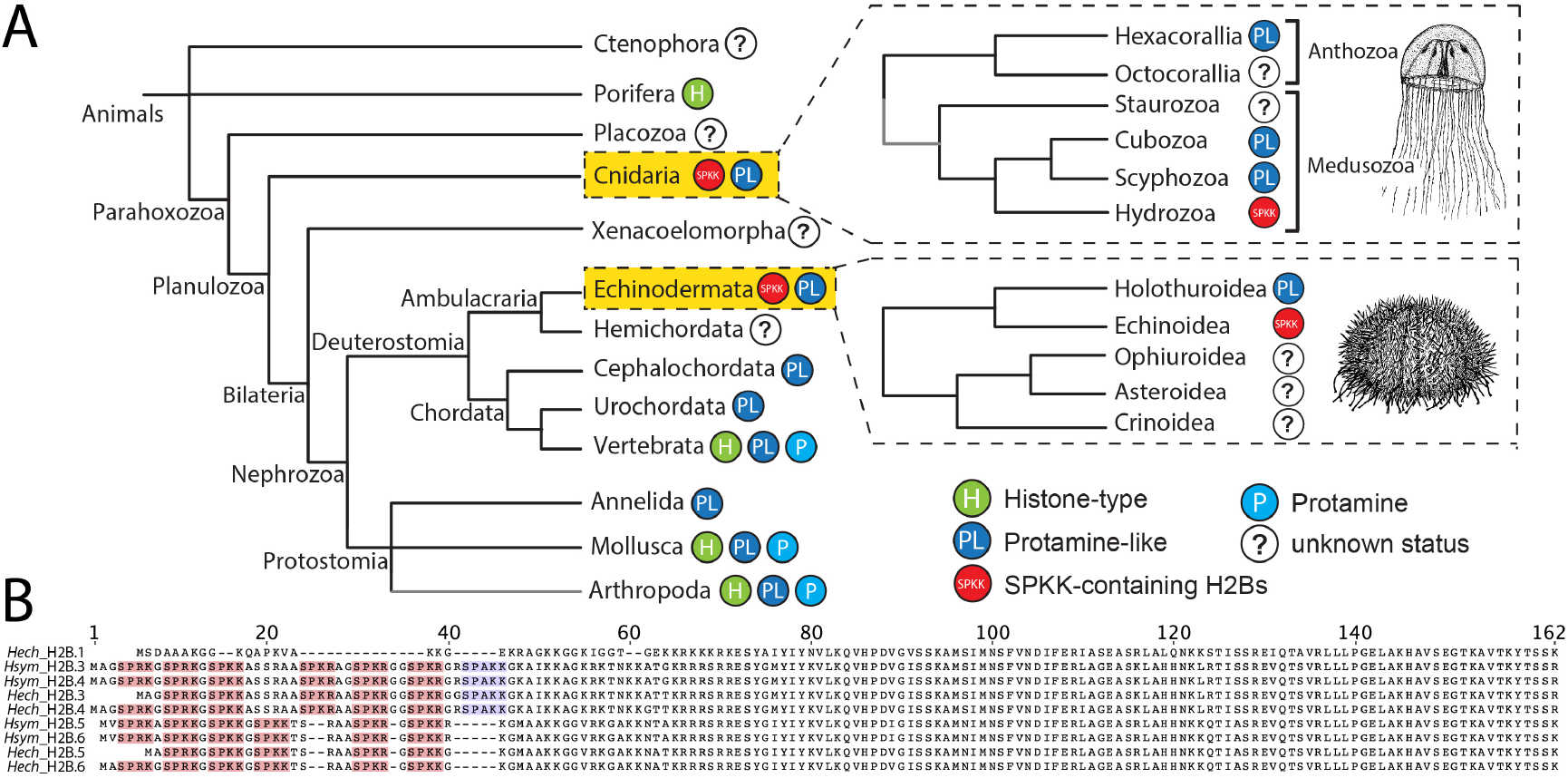
The evolution of N-terminal SPKK-related motif-containing H2B histones. (A) Cladogram depicting the distribution of SNBPs in animals. (B) Protein alignment showing *H. echinata* and *H. symbiolongicarpus* canonical and SP[K/R][K/R]-containing H2Bs.

We have previously shown that spH2Bs are expressed in male gonads of the hydrozoan cnidarian *Hydractinia echinata* (Török et al., 2016). Here, we characterise the *Hydractinia* spH2Bs biochemically and functionally using the sibling species *H. symbiolongicarpus*. We complement this with *in vitro* observations of the properties of mononucleosomes and oligonucleosomal arrays containing recombinant *Hydractinia* spH2Bs. Our results show that spH2Bs inhibit transcription and stabilize chromatin without contributing significantly to its compaction. These characteristics are consistent with a function to facilitate fertilization in these broadcast-spawning animals.

## Results

### spH2Bs are co-expressed and replace H2B.1 during spermatogenesis but do not significantly increase chromatin compaction

*Hydractinia echinata* and *H. symbiolongicarpus* both possess several hundred, tandemly repeated copies of a single canonical H2B-encoding gene (*H2B.1*) but only four non-canonical sperm-specific H2B variant (*H2B.3, H2B.4, H2B.5, H2B.6*; Fig. 1B); the latter containing 5-7 N-terminal SP[K/R][K/R] repeats (Török et al., 2016). The H2B.3 and H2B.4 pair, and the H2B.5 and H2B.6 pair, are each identical in protein sequence but differ slightly at the nucleotide level and are encoded at unique chromosomal loci (Török et al., 2016). We refer to all 4 variants collectively as spH2Bs, except where a specific member is investigated. For *in vitro* experiments, we investigated the properties of *H. echinata* H2B.3 and H2B.6 variants as representatives of *Hydractinia* spH2B pairs and compared them with canonical H2B.1.

We used fluorescence in situ hybridization (FISH) to confirm that the expression patterns of spH2Bs in *H. symbiolongicarpus* were equivalent to the previously published *H. echinata* expression. As reported (Török et al., 2016), the high nucleotide sequence similarity between *H2B.3* and *H2B.4*, and between *H2B.5* and *H2B.6*, precludes the design of gene-specific cRNA probes so we treated each pair as a combined case. We observed that *H2B.3/4* and *H2B.5/6* were co-expressed in sperm progenitor cells within male gonads of *H. symbiolongicarpus* (Fig. 2A) while the expression of *H2B.1* was concomitantly downregulated (Török et al., 2016), consistent with its replacement by spH2Bs during spermatogenesis.

**Figure 2:**
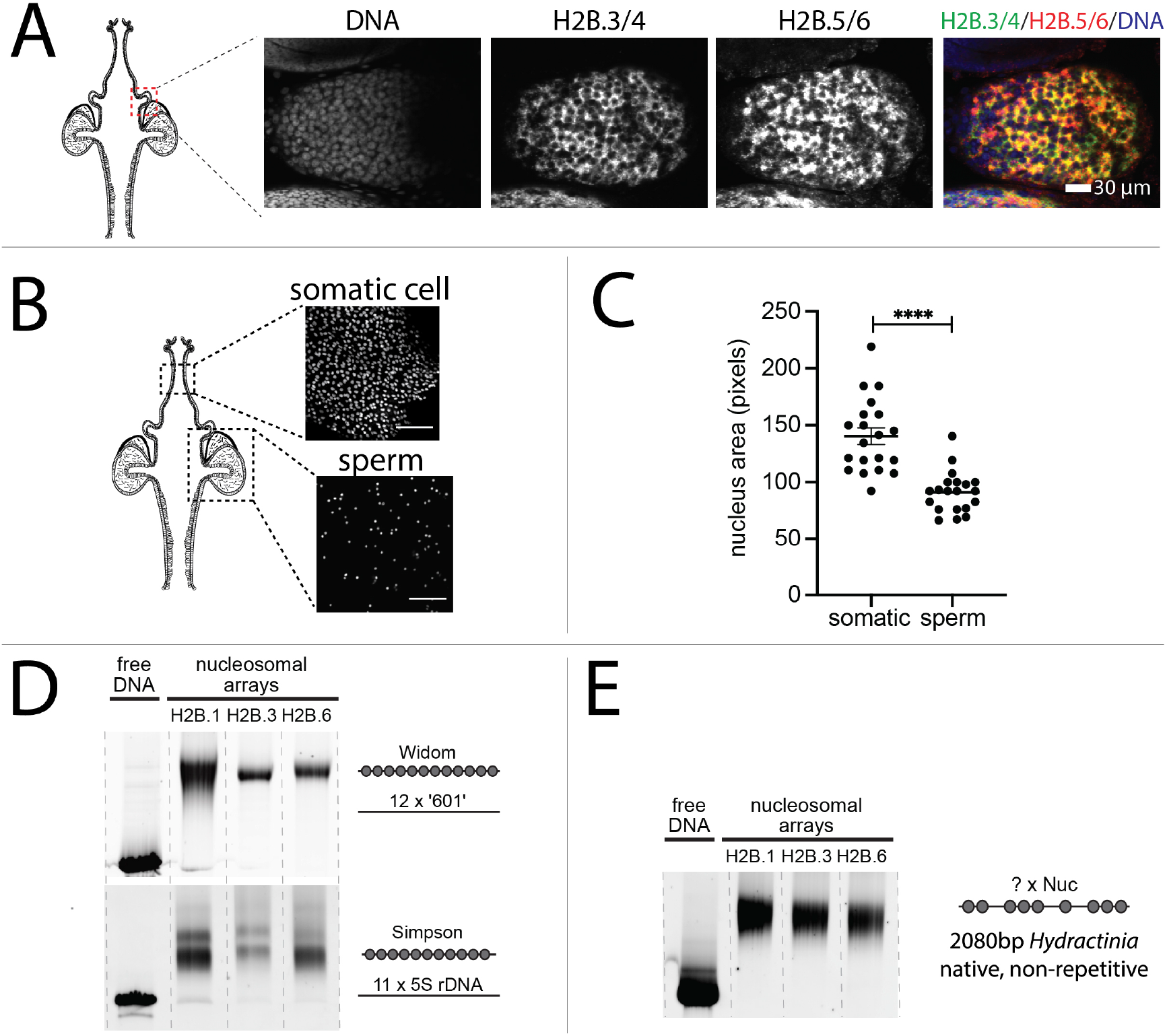
spH2Bs are co-expressed, replace H2B.1, but do not enhance chromatin compaction. (A) Expression of *H2B.3*/*H2B.4* and *H2B.5*/*H2B.6* in sperm progenitor cells visualised by RNA FISH. (B) Hoechst staining of DNA in diploid somatic cells and mature haploid sperm cells for nuclear area measurements. Scale bar 10 μm. (C) Distribution of nuclear areas for cell types in B (n=20). (D) Native PAGE of nucleosome arrays assembled on 12x 177 bp 601 repeat (‘Widom’) and 11x 208 bp 5S rDNA repeat (‘Simpson’) using histone octamers containing either H2B.1, H2B.3 or H2B.6. (E) Native PAGE of a 2080 bp non-repetitive *Hydractinia* DNA using the same octamers as in (E).

To test whether spH2Bs affect overall genome compaction relative to H2B.1, we measured the nuclear dimensions of sperm cells relative to somatic cells in *H. symbiolongicarpus*. Nuclei of male sexual polyps were stained with Hoechst and confocal images were taken of somatic cells and of cells in the late stage of spermatogenesis (Fig 2B). Then, the cross-section areas of multiple individual nuclei were calculated. Nuclei of diploid somatic cells had a median area that was 50% larger than haploid mature sperm nuclei (Fig 2C). This is consistent with little or no volume reduction of chromatin involving spH2Bs comparing with 6-8-fold volume reduction in protamine-compacted sperm chromatin (Fuentes-Mascorro et al., 2000).

To further investigate the DNA compaction by spH2Bs, we assembled nucleosomal arrays from *Hydractinia* histones. Oligonucleosome arrays containing 12 repeats of the well-characterised Widom 601 strong nucleosome positioning sequence (Dorigo et al., 2003; Lowary and Widom, 1998) were assembled with recombinant *Hydractinia* histone octamers containing either H2B.1, H2B.3, or H2B.6. Native PAGE analysis of these nucleosomal arrays did not show any significant difference in the electrophoretic mobility (Fig. 2D). To ensure that this was not an artifact of the 177 bp Widom 601 sequence repeat, we also prepared equivalent oligonucleosome arrays on the 12 x 208 bp repeat Simpson 5S sequence (Simpson et al., 1985) and on a native non-repetitive 2080 bp *Hydractinia* genomic AT-rich DNA sequence with equivalent results (Fig. 2E). We conclude that spH2Bs do not lead to significantly enhanced chromatin compaction compared to H2B.1 *in vitro* or *in vivo*.

### SP[K/R][K/R] motifs stabilise chromatin structure and restrict chromatin accessibility

To observe the effect of spH2Bs on nucleosome stability, we assembled mononucleosomes using recombinant *Hydractinia* histone octamers containing H2B.1, H2B.3, or H2B.6 separately as mononucleosomes and measured their thermal stability (Taguchi et al., 2014). DNA was fully released from all mononucleosomes at 88-90°C, and nucleosomes containing canonical H2B.1 showed a maximum of H2A-H2B dimer dissociation at 70°C (Fig. 3A), consistent with other metazoan canonical H2B-containing nucleosomes (Taguchi et al., 2014). In contrast, nucleosomes incorporating either H2B.3 or H2B.6 showed more progressive thermal dissociation of H2A-H2B dimers, reaching a maximum at 78°C (Fig. 3A).

**Figure 3:**
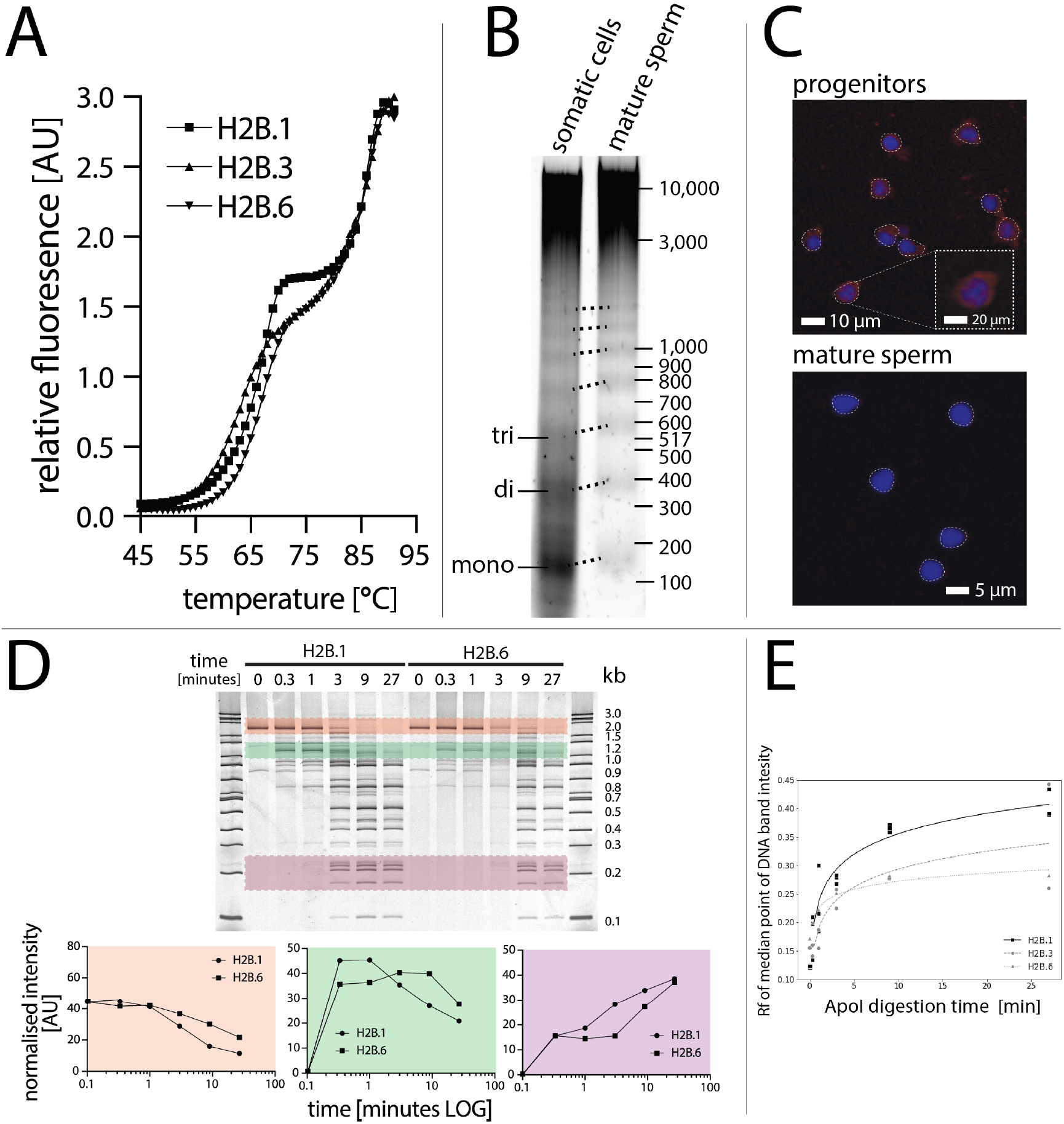
spH2Bs stabilise chromatin structure and restrict chromatin accessibility *in vivo* and *in vitro*. (A) Stability of mononucleosomes containing recombinant H2B.1, H2B.3, or H2B.6 measured by temperature dependence of fluorescent dye binding *in vitro*. (B) Difference in nucleosomal repeat for somatic and mature sperm cells after MNase digestion *in vivo*. (C) Mature sperm are inaccessible to Tn5 integrase (red) comparing to progenitors. Nuclei were counterstained with Hoechst (blue). (D) Native PAGE of equimolar recombinant H2B.1 or H2B.6-containing nucleosome arrays on the 2080 bp non-repetitive *Hydractinia Piwi1* 5’ non-coding region DNA digested by ApoI restriction enzyme. Band intensities for 3 representative DNA bands are shown (pink, red and purple). (E) Quantification of native PAGE migration (Rf) for median point of DNA band intensity for ApoI digestion of H2B.1, H2B.3 or H2B.6-containing nucleosomal arrays from 4 replicate native PAGE, showing reduced accessibility of H2B.6 nucleosomes *in vitro*.

To probe the impact of spH2Bs on chromatin accessibility *in vivo*, we performed nuclease accessibility assays and observed an increase in nucleosomal repeat lengths after micrococcal nuclease (MNase) digestion of mature sperm cells compared to progenitors, consistent with a reduced accessibility of linker DNA between spH2B-containing nucleosomes (Green and Poccia, 1988; Hill and Thomas, 1990) (Fig. 3B). We then performed ATAC-see on these nuclei and observed a strong signal from sperm progenitors whereas DNA was inaccessible to Tn5 transposase in mature sperm (Fig. 3C). This demonstrates that sperm chromatin has reduced accessibility *in vivo*.

Finally, we performed parallel *in vitro* ApoI restriction enzyme digestion on oligonucleosomal arrays assembled *in vitro* on a native 2080 bp non-repetitive *Hydractinia* DNA fragment with H2B.1, H2B.3, or H2B.6 containing histone octamers. ApoI digestion at 8 recognition sites along the DNA was quantitatively slower for H2B.3 or H2B.6-containing arrays than for H2B.1-containing arrays (Fig. 3D and 3E), in agreement with the reduction of accessibility seen *in vivo*.

### spH2Bs cause a transcription block and cell cycle arrest in the absence of H2B.1

To test the functional effects of spH2B incorporation in chromatin we injected *in vitro* transcribed mRNA encoding the full-length sequences of *H2B.3, H2B.4, H2B.5*, and *H2B.6* fused to GFP together into one blastomere of two-cell stage embryos and compared this to injections of mRNA encoding GFP alone as a control. All embryos showed GFP fluorescence 6 hours after injection in 50% of cells (Fig. 4A). Animals injected with the combined spH2B-GFP fusion mRNAs exhibited nuclear fluorescence whereas control *GFP* alone was cytoplasmic (Fig. S1). Embryos developed into planula larvae within 2-3 days, similar to untreated *Hydractinia* embryos.

**Figure 4:**
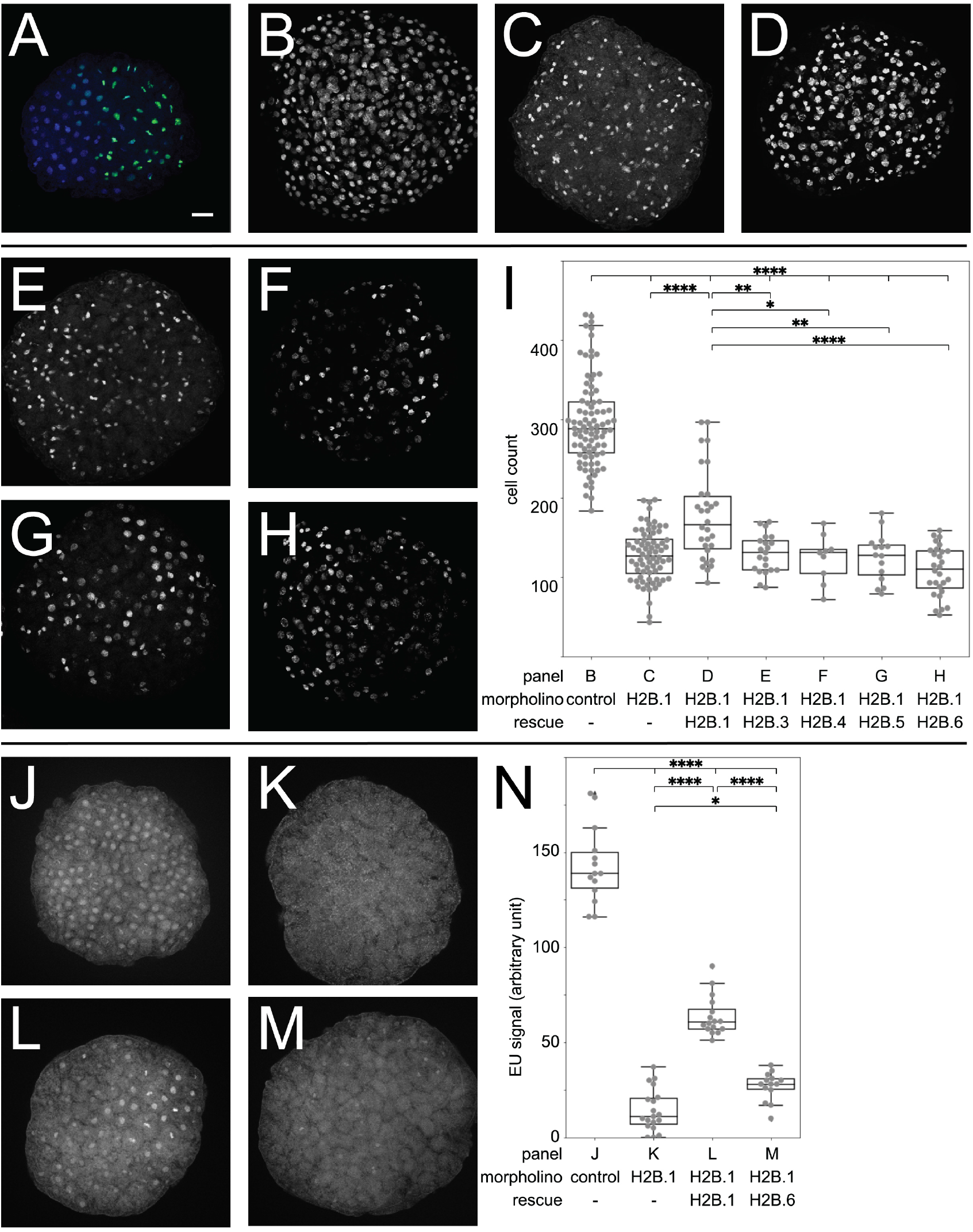
spH2Bs cause a transcription block and cell cycle arrest in the absence of H2B.1. (A) GFP fluorescence 7 hours after injection of GFP control mRNA into embryos. (B-H) Embryos stained with Hoechst after injection with (B) control morpholino, (C) H2B.1 translation-blocking morpholino, (D) H2B.1-blocking morpholino and morpholino-resistant H2B.1 mRNA, (E) H2B.1-blocking morpholino and H2B.3 mRNA, (F) H2B.1-blocking morpholino and H2B.4 mRNA, (G) H2B.1-blocking morpholino and H2B.5 mRNA, (H) H2B.1-blocking morpholino an H2B.6 mRNA. (I) Distribution of cell counts in embryos from panels B-H. (J) EU incorporation in embryo injected with control morpholino. (K) EU incorporation in embryo injected with H2B.1-blocking morpholino. (L) EU incorporation in embryo injected with H2B.1-blocking morpholino and morpholino-resistant H2B.1 mRNA. EU incorporation in embryo injected with H2B.1-blocking morpholino and H2B.6 mRNA. N. Quantification of EU signals in embryos from panels J-M. The scale bars equal 20 μm.

We hypothesized that the injected mRNAs encoding spH2Bs would be outnumbered by endogenous *H2B.1* transcripts because there are some 700 copies of the canonical *H2B.1* gene in the *Hydractinia* haploid genome but only single copies of each spH2B variant (Török et al., 2016). Therefore, we co-injected an *H2B.1*-specific translation blocking morpholino together with the mRNAs for the four spH2B variants (Fig. S2). Control morpholinos had no effect on the embryos (Fig. 4B) and injecting the morpholino alone led to cell cycle arrest and death of all embryos within a few cell cycles (Fig. 4C). This could be partially rescued by co-injection of morpholino-resistant *H2B.1* encoding mRNA (Fig. 4D and Fig. S2), confirming the specificity of the morpholino.

We then injected *H2B.3, H2B.4, H2B.5*, and *H2B.6* mRNAs individually along with the *H2B.1*-specific morpholino. This caused cell cycle arrest equivalent to the injection of the morpholino alone, showing that spH2Bs cannot substitute for H2B.1 (Fig. 4E-I). Finally, we used ethynyl uridine (EU) incorporation to quantitate nascent RNA levels and found them to be significantly lower in *H2B.6* mRNA-injected embryos than in embryos co-injected with the *H2B.1* rescue mRNA in morphants (Fig. 4J-N).

Therefore, the mRNA injection experiments are consistent with spH2B incorporation in chromatin in place of H2B.1 leading to a generalised transcriptional block.

## Discussion

Sperm-specific histones H2B.3, H2B.4, H2B.5 and H2B.6 containing N-terminal SP[K/R][K/R] repeats increased chromatin stability and reduced DNA accessibility compared with chromatin assembled with canonical H2B.1 both *in vivo* and *in vitro*. Our observations are consistent with transcriptional silencing resulting from networks of linker DNA binding of spH2Bs in sperm, in line with earlier investigations with structurally similar H2B variants in sea urchin sperm. However, given the suggested ease of evolving SPKK-related motifs (Malik et al., 2002) and the fact that spH2Bs are only known in two unrelated taxa, we suggest that hydrozoan and echinoid spH2Bs evolved convergently.

Protamine-type SNBPs generate highly compact nuclei and this has been suggested to improved sperm motility (Champroux et al., 2016; Levitan and Petersen, 1995). In contrast, *Hydractinia* spH2Bs do not appear to increase chromatin density. *Hydractinia* colonies are dioecious, growing individually on the surface of hermit crab shells. Their spawning is light induced but the distance to a potential mate depends on the behaviour of their host hermit crabs. Eggs are near neutrally buoyant upon spawning, sinking only very slowly to the bottom in calm water (Video S1) and remaining suspended under turbulent conditions. This feature probably prevents egg loss in the sediment, but fertilization would be highly unlikely unless sperm have comparable buoyancy. Upon fertilization, zygotes become strongly negatively buoyant (Video S2; Fig. S3), probably to protect the embryo from predation in the water column and allow the larvae to attach to benthic hermit crab shells. Since chromatin contributes a major part of total sperm biomolecule composition, we speculate that by maintaining reduced accessibility without increasing compaction, spH2Bs could contribute to neutral *Hydractinia* sperm buoyancy. Less dense sperm would increase fertilization likelihood and could be a selective pressure that drove the evolution of hydrozoan spH2Bs.

## Material and Methods

### Animal culture

*Hydractinia* colonies were cultured in artificial seawater at 18°C under 14h:10h light– dark regimes and fed *Artemia franciscana* nauplii four times a week, and ground oyster once per week. Spawning is light induced, occurring synchronously in both sexes (Frank et al., 2020). Fertilised eggs were collected for microinjection and developed at room temperature into planula larvae within 3 days. Polyps were harvested from mature colonies.

### Microinjection

Injection needles were prepared from glass capillaries (Narishige GD-1 1×90 mm) using a microneedle puller (Narishige; cat no.: PN-31) with settings of heat 560, pull 70, velocity 75, time 150. Microinjection was carried out immediately after fertilisation in 1-cell stage embryos. Embryos were placed in a small Petri dish lid with a 200 μm plankton net attached to avoid any movement. Injections were completed before the first cell division.

### Genomic DNA extractions

As previously described (Török et al., 2016).

### Micrococcal nuclease (MNase) assay

As previously described (Török et al., 2016).

### Fluorescent in situ hybridisation (FISH)

As previously described (Török et al., 2016).

### Capped and polyadenylated mRNA synthesis

For mRNA synthesis the desired fragment including 100 bp 5’-UTR and the coding sequence of the histone gene was amplified by PCR from genomic DNA. Linker sequences and GFP tags were added to the construct by Gibson assembly. The assembled fragment was re-amplified by PCR, with primers containing T7 promoter necessary for RNA synthesis. HiScribe T7 ARCA mRNA kit (NEB) was used for mRNA synthesis with 10 μl of 2x ARCA/NTP mix, 1 μg of template DNA, 2 μl of T7 RNA Polymerase mix and nuclease-free water to a final volume of 20 μl. Reactions were gently mixed and incubated at 37°C for 1-2 hours. 2 μl DNaseI was added to the mixtures and incubated for a further 15 min. PolyA tailing was carried out in a total volume of 100 μl with 20 μl of IVT reaction (from above), 65 μl of nuclease-free water, 10 μl of 10x Poly(A) Polymerase Reaction Buffer and 5 μl of Poly(A) polymerase. Reactions were mixed and incubated at 37°C for 30 min. In order to purify mRNA 0.5 volume of LiCl solution (7.5 M LiCl, 10 mM EDTA) was added to each sample and incubated at −20°C for 30 min. Tubes were centrifuged at 4°C for 15 min at full speed. Supernatant was removed and pellets were rinsed with 500 μl cold ethanol. Tubes were centrifuged again at 4°C for 10 min at full speed. Ethanol and residual liquid was carefully removed using a sharp tip then heated to 65°C for 5-10 min to completely dissolve RNA. Following this, mRNA concentrations were measured and the tubes were stored at −80°C. mRNA could be visualised and analysed on formaldehyde denaturing gels.

### EdU staining of S-phase cells

EdU visualisation was performed using a Click-iT EdU Alexa Fluor 488 Imaging kit (ThermoFisher Scientific). The solutions were prepared according to the manufacturer’s instructions. Fluorescence excitation and emission maxima for Alexa Fluor 488 were 495 and 519 nm respectively. For EdU staining animals were incubated in EdU solution for 30 min at a concentration of 150 μM. Following incubation FISH was performed as described above when it was necessary. Animals were then fixed in 4% PFA in 100 mM HEPES pH 7.5, 4 mM MgSO4, 140 mM NaCl overnight at 4°C. Following this, animals were washed twice in 3% BSA/PBSTx (0.1%) for 15 min each. The sample was permeabilised by incubation in 0.5% PBSTx for 1 hour. Animals were then washed twice for 10 min in 3% BSA/PBSTx (0.1%).

Staining cocktail containing 430 μl of 1x reaction buffer, 20 μl CuSO_4_, 1.2 μl Alexa Fluor 488 azide and 50 μl of reaction buffer additive in a total volume of 0.5 ml was prepared. The reaction cocktail was then added to the animals to cover them entirely and incubated in the dark for 30 min at RT then washed four times in 3% BSA/PBS Triton (0.1%) for 15 min each. Nuclei were stained in 10 ng μl^−1^ Hoechst 33258 (Sigma B2883) in PBSTx for 15 min followed by one wash in PBSTx for 10 min. Samples were mounted on microscopic glass slides using Fluoroshield mounting medium (Sigma F4680), sealed with nail polish and stored at −20°C. Images were taken on Olympus FV1000 inverted confocal microscope.

### EU staining to detect nascent RNA

EU visualisation was performed using a Click-iT RNA Imaging Kit (ThermoFisher Scientific). The solutions were prepared according to the manufacturer’s instructions. For EU staining embryos were incubated in EU solution for 1 hour at a final concentration of 0.5 mM and fixed in 4% PFA in HEPES for 1 hour at RT. Embryos were then washed in PBS for 5 min followed by quick washes in increasing concentrations of methanol. Animals were first placed in 25% methanol v/v in PBS followed sequentially by 50%, 75% and 100% methanol. The embryos were then rehydrated by washing in decreasing concentrations of MeOH. First in 75% MeOH, then in 50% methanol and finally in 25% methanol for a few seconds each. Embryos were washed once again in PBS for 5 min. To permeabilise the samples, the embryos were incubated in 0.5% TritonX-100 in PBS for 15 min at RT. The permeabilisation buffer was removed and the embryos were washed once with PBS for 5 min. Staining cocktail was prepared to a total volume of 250 μl containing 214 μl of 1x Click-iT RNA reaction buffer, 10 μl CuSO4, 1 μl Alexa Fluor 488 azide and 25 μl Click-iT reaction buffer additive. The reaction cocktail was added to the embryos to cover them entirely. The animals were incubated for 30 min at RT, protected from light. Following incubation animals were washed four times in PBS for 15 min each. Nuclei were stained in 10 mg ml^−1^ DAPI in PBS for 15 min. Animals were washed in 0.3 % TritonX-100 in PBS for 10 min. Samples were then mounted on microscopic glass slides using Fluoroshield mounting medium (Sigma F4680), sealed with nail polish and stored at −20°C. Images were taken on Olympus FV1000 inverted confocal microscope.

### Microscopy

An Olympus SZX7 stereomicroscope was used for animal observation, colony maintenance and injection. Fluorescence microscopy was carried out using an Olympus BX51 compound microscope. Images were processed with the Olympus CellD software package. Confocal microscopy was carried out on the Olympus Fluoview 1000 software with an inverted IX71 microscope. In order to reduce noise the Kalman accumulation function was applied. Images were then acquired and saved. In order to observe the desired multiple sections the Z-position was also adjusted. The stacks of sections could then be projected together. Image analysis was carried out using ImageJ Software.

### Morpholino mediated knockdown

Morpholino oligos were designed around the start codon of the gene of interest using Gene Tools software (http://www.gene-tools.com) (Fig. S1A). Morpholino oligos were diluted to the recommended concentrations and injected into 1-cell stage embryos. Control morpholino sequence: CCTCTTACCTCAgTTACAATTTATA. H2B.1 morpholino sequence: GCTGCTGCGTCAGACATGGTTAAAT.

### Assay of transposase-accessible chromatin with visualisation (ATAC-see)

Hyperactive Tn5 transposase was produced and purified as described (Picelli et al., 2014). High-quality, tagmentation-ready Tn5 transposase was then loaded with DNA adaptors that selectively bind to accessible chromatin only (Buenrostro et al., 2013). Tn5 transposome assembly was carried out by resuspending Tn5ME-AATTO590, Tn5ME-B-ATTO590 and Tn5MErev oligos in nuclease free water to a final concentration of 100 μM. Equimolar amounts of Tn5MErev/Tn5ME-A-ATTO590 and Tn5MErev/Tn5ME-B-ATTO590 were mixed in separate 200 μl PCR tubes and incubated at 95°C for 5 min then cooled down slowly by turning off the thermocycler. The assembly of transposase solution was carried out in dark using 0.25 volumes of Tn5MErev/Tn5ME-A-ATTO590 + Tn5MErev/Tn5ME-B-ATTO590 at 50 μM final concentration, 0.4 volume of 100% sterile glycerol, 0.12 volume of sterile filtered 2x dialysis buffer (100 mM HEPES–KOH at pH 7.2, 0.2 M NaCl, 0.2 mM EDTA, 2 mM DTT, 0.2% Triton X100, 20% glycerol), 0.1 volume of 50 μM SL–Tn5 and 0.13 volumes of water. Reagents were gently mixed, and incubated at RT for 1 h, allowing the oligos to anneal to Tn5. Transposase solution was stored at −20°C. Slide preparation and fixation was carried outby fixing *Hydractinia* sperm cells and pronase-treated somatic cells with 1% formaldehyde for 10 min at RT. Sperm cells were fixed on glass coverslips using Cytospin, however this could not be performed with somatic cells as it damaged their structure. ATAC-see staining was carried out by premeabilising with lysis buffer (10 mM Tris–Cl, pH 7.4, 10 mM NaCl, 3 mM MgCl2, 0.01% Igepal CA-630) for 10 min at RT. The samples were rinsed with 1x PBS twice and placed in a humid chamber box at 37°C. The transposase mixture solution (25 μl 2x TD buffer, final concentration of 100 nM Tn5ATTO-59ON in a volume of 50 μl) was placed on the cells and incubated for 30 min at 37°C. The slides were washed with 1x PBS containing 0.01% SDS and 50 mM EDTA for 15 min three times at 55°C. Nuclei were stained in 10 ng μl^−1^ Hoechst 33258 (Sigma B2883) then mounted using Fluoroshield medium (Sigma F4680) for confocal imaging.

### Recombinant histone expression

All histone genes except H4 were amplified from *Hydractinia* genomic DNA using degenerate primers and subcloned into pET3a. The H4 coding sequence was optimized and synthesized in pD451 (Suppl. File S1). *E. coli* Rosetta2 pLysS or Star pRIL cells chosen for optimal expression were transformed with expression plasmids and plated on LB agar containing appropriate antibiotics then grown overnight at 37°C. Colonies were transferred to 1 L of 2YT media supplemented with antibiotics and grown until OD600 0.6-0.8. Expression was induced by the addition of IPTG to a final concentration of 0.4 mM. Cultures were grown for a further 4 h at 37°C and 180 rpm. Cells were then harvested by centrifugation at 5,000 G for 15 min and cell pellets were stored at −20°C.

### Recombinant histone purification

The cell pellet containing histone inclusion bodies were allowed to thaw and resuspended in 30 ml Histone Wash Buffer (50 mM Tris-HCl pH7.5, 100 mM NaCl, 5 mM β-mercaptoethanol, 1 mM benzamidine hydrochloride). Cell suspensions were sonicated in a Branson Sonifier 250 with 12 mm tip on ice for 1 min at 40% amplitude with 5 s on pulses and 10 s off pulses, and the lysate was centrifuged at 30,000 G for 15 min. The resulting pellet was washed several times with Histone Wash Buffer, resuspended in 0.5 ml DMSO and incubated at RT for 30 min. The inclusion bodies were further solubilised by the addition of 5 ml of Unfolding Buffer (20 mM Tris-HCl pH 7.5, 7 M Guanidine HCl, 10 mM DTT) and incubated on rollers for 1 h at RT. Following this, Histone Purification Buffer A (50 mM Tris-HCl pH 7.5, 7 M Urea, 1 mM EDTA) was added to a final volume of 20 ml and centrifuged at 35,000 G for 20 min. The resulting supernatant was centrifuged for 20 min at 35,000 G. This final supernatant was then added to 55 ml of Histone Purification Buffer A. The histone solution was passed through a PVDF filter with 0.2 μm pore size attached to a vacuum bottle. Samples were then loaded onto a 5 ml SP Sepharose HP column using an FPLC system. Histones were eluted with a linear gradient of increasing Histone Purification Buffer B (50 mM Tris-HCl pH 7.5, 7 M Urea, 1 mM EDTA, 2 M NaCl).

### SDS polyacrylamide gel electrophoresis (SDS-PAGE)

SDS polyacrylamide gels were made to a final acrylamide concentration of 15% acrylamide with 1:37.5 bis-acrylamide. Samples were resuspended in the presence of 1x protein gel loading buffer and heated to 95°C for 5 min prior to loading. Gel electrophoresis was carried out by applying a constant 100 V in 1x TG buffer (30 g L^−1^ Tris, 144 g L^−1^ glycine). Protein bands were visualised by Coomassie blue staining.

### Histone octamer refolding by salt dialysis

Equal quantities of lyophilised histones were resuspended in Unfolding Buffer to a final concentration of 2 mg ml^−1^. Denaturing resuspension was carried out at RT on a roller for 1 h then centrifugation at 18,000 G for 10 min. The concentration of each histone supernatant was determined using calculated extinction coefficients (Expasy ProtParam). The four histones were mixed in equimolar ratios and the volume adjusted to a final protein concentration of 1 mg ml^−1^. This mixture was placed in an 8,000 MWCO dialysis bag and twice dialysed against 600 ml of Refolding Buffer (10 mM Tris-HCl pH 7.5, 2 M NaCl, 1 mM EDTA, 5 mM β-mercaptoethanol) at 4°C for 3 h each followed by a third overnight dialysis. After concentration to 1 ml using a Millipore Ultrafree 10K MWCO concentrator, histone octamers were purified by size exclusion chromatography using a Superdex 200 column equilibrated in Refolding Buffer. Fractions containing all 4 histones at equimolar ratios were confirmed by SDS PAGE and pooled then concentrated to approx. 10 μM using a Millipore Ultrafree 10K MWCO concentrator.

### DNA preparation by plasmid digestion

Plasmids containing 12x 177 bp 601 (Dorigo et al., 2003) and pCL3 containing 11x 208 bp 5S rDNA DNA repeats (Logie and Peterson, 1997) were transformed into *E. coli* TOP10 cells then grown in large scale LB media cultures with appropriate antibiotics. Plasmids were extracted from harvested cells by using ThermoFisher GeneJet plasmid DNA purification kits.

The 601 repeat insert was isolated by digesting purified plasmid at 37°C overnight using 0.34 NEB units μg^−1^ of EcoRV. The 5S repeat insert was isolated by digesting pCL3 with EcoRV, BglI and BstXI at 37°C overnight using 0.34 NEB units μg^−1^ of each enzyme. The 2.1 kbp and 2.3 kbp inserts were isolated from smaller plasmid fragments after electrophoresis in 0.7% agarose gels run in 1x TAE buffer (40 mM Tris-acetate pH 8.0, 1 mM EDTA) using a ThermoFisher GeneJet gel extraction kit.

Finally, DNA fragments were loaded onto a 5 mL MonoQ ion exchange column and eluted by a linear gradient of DNA binding buffer B (20 mM Tris-HCl pH7.5, 0.1 mM EDTA, 2 M NaCl) over 5 column volumes. The desired peak fractions were identified by 1% agarose gel electrophoresis, then ethanol precipitated and the pellet was resuspended in 10 mM Tris-HCl pH 7.5.

### DNA preparation by PCR

Two 96 well plates of identical PCR reactions with 400 pg μL^−1^ template plasmid and 400 μM dNTPs in Taq buffer (10 mM Tris-HCl pH 9.0, 50 mM KCl, 1.5 mM MgCl_2_, 0.1% Triton X-100) with were amplified using Taq polymerase with 800 nM primers. Template plasmids containing a 2080 bp non-repetitive region of the *H. echinata* Piwi1 promoter (Suppl. File S1; F- CAGATGATCCGCAGACAATAG, R-AAATGTAATGAAAATTTTCGTAATTA) or the Widom ‘601’ sequence (F-CTGCAGAAGCTTGGTCCC, R-ACAGGATGTATATATCTG) were used for the 2080 bp and 147 bp fragments, respectively. PCR wells were pooled and ethanol precipitated by adding 0.1 volumes of 3 M sodium acetate pH 5.2 and 2.5 volumes of absolute ethanol then centrifuging at 11,000 G at 4°C for 30 min. The DNA pellet was resuspended in 2 ml of DNA binding buffer A (20 mM Tris-HCl pH 7.5, 0.1 mM EDTA). Finally, DNA was purified by ion exchange chromatography as described above.

### Nucleosome assembly by salt dialysis

To assemble 250 pmol of nucleosomes in a volume of 80 μl, purified octamer was mixed with the appropriate DNA in an equimolar ratio of nucleosome sites in a solution containing a final concentration of 20 mM Tris-HCl pH 7.5 and 2 M NaCl. The reaction mixture was transferred into a custom-made miniature dialysis block pre-equilibrated at 4°C with 8,000 MWCO dialysis membranes. Samples were dialysed against solutions containing 10 mM Tris-HCl pH 7.5 and 1 mM EDTA with consecutively 1.4 M, 1.2 M, 0.8 M and 0.6 M NaCl for at least 2 h each, and then finally against 10 mM Tris-HCl pH 7.5 overnight. Nucleosomal arrays were visualised by mixed agarose-polyacrylamide native gel electrophoresis, and mononucleosomes were visualised by native PAGE.

### Mixed agarose-polyacrylamide native gel electrophoresis

Gels comprising 1% agarose and 2% acrylamide with 1:37.5 bisacrylamide in 0.2x TBE were heated to dissolve agarose, where 1x TBE is 89 mM Tris pH 7.5, 89 mM borate, 2 mM EDTA. The solution was allowed to cool to ~40°C before adding 0.1% TEMED and ammonium persulphate to 0.1% then pouring into pre-warmed 16 cm x 20 cm glass plates with 1 mm spacers. Polymerised gels were pre-equilibrated for 145 min at 250 V in 0.2x TBE Buffer. Samples were mixed with 0.7 volumes of 10 mM Tris-HCl pH 7.5, 6% sucrose. Electrophoresis was performed under the same conditions as pre-equilibration.

### Nucleosome array digestion by ApoI

Nucleosome arrays were digested with ApoI restriction enzyme by mixing 15 pmol assembled nucleosomal array in 50 μl volume with 1x NEB CutSmart buffer and 7.5 units of restriction enzyme at 37°C. 10 μL samples were withdrawn at each time point and mixed rapidly with 33% phenol and 33% chloroform to stop the reaction. The aqueous phase was separated by centrifugation then ethanol precipitated, and the DNA pellet was resuspended in 1x DNA loading dye. Digestion was visualised by native PAGE.

### Native PAGE

A 6% acrylamide with 1:37.5 bisacrylamide native polyacrylamide gel was cast and left for 1 h at RT for complete polymerization. The gel was thermally equilibrated for 1 h at 4°C before being pre-run for 3 h at 250 V and 4°C in 0.2x TBE. 4 pmol of nucleosome or DNA in 5% sucrose was loaded per lane in native PAGE and the gel was run for a further 3 h at 250 V at 4°C in 0.2x TBE.

### Imaging of native polyacrylamide gels

Gels with unlabelled DNA were stained with 10 ng ml^−1^ ethidium bromide for 15 min then scanned using a Fuji FLA5100 or BioRad Pharos FX fluorescent imager with the appropriate laser and filter settings for Cy3, Cy5 or ethidium bromide.

### Thermal stability assay by fluorescent dye binding

A 20 μL reaction mix was prepared by mixing 36 pmol of nucleosomes assembled on 147 bp Widom 601 DNA with freshly diluted 5x SyPro Orange with 0.1 mM NaCl, 20 mM Tris-Cl, pH 7.5 in a 96 well MicroAmp Fast optical plate. Thermal unfolding analysed on StepOne Plus Thermal Cycler as described in (Taguchi et al., 2014).

### Cell counting and image analysis

The Automated Counting feature of ImageJ/Fiji (https://fiji.sc/) was used for cell counting. For EdU positive nuclei counting, confocal Z-stacks were projected. Color images (RGB) were first converted into greyscale before proceeding through the following settings: ‘Edit – Options – Conversations’ to ‘scale when converting.’ Then images were convert to greyscale using ‘Image - Type - 16-bit’. All structures to be counted were highlighted according to the following settings: ‘Image - Adjust – Threshold’. The sliders were then used to highlight structures and settings were applied to create a binary version of the image with pixel intensities black = 0 and white = 255. When particles overlapped the ‘Process - Binary – Watershed’ function was applied to separate then by adding a 1 pixel thick line. Following this, the ‘Analyze - Analyze Particles’ function was used. In order to avoid counting noise, particle size was adjusted excluding noise-derived particles. The outlines of cell nuclei were then appearing together with a summary table showing the number of cells counted. These results were then used for statistical analysis. Cell counting experiments are shown as data from 2-3 independent technical replicates, with 15-20 biological replicates in each group. Data was normalised and p values were calculated using T-test in the SPSS software package.

### Analysis of nuclear areas

Nuclei of intact male sexual polyps were stained by Hoechst and confocal images from early to late stages of spermatogenesis were taken, along with epithelial somatic cells from the surface of the body column with the same magnifications and settings. The perimeter of 100 nuclei in an optical section of each sample was defined manually and the pixels counted as a measurement of area.

### Analysis of native gel migration

Individual lanes from immediately below the well to the smallest DNA band were cropped from TIFF images and read as pixelwise arrays then summed by row. Background was estimated from the region of 97% migration in image and subtracted, then the row-wise cumulative intensity was calculated and normalised to the maximum. The relative migration of the 50^th^ percentile for the cumulative signal was extracted as the Rf of median DNA band intensity versus ApoI digestion time. Time series data for histone octamers containing each histone variant (H2B.1 n=3, H2B.3 n=2, H2B.6 n=1) were fitted to a log normal function using curve_fit from the optimize module of SciPy. All data was plotted using seaborn with appropriate matplotlib parameters.

## Declarations

### Ethics approval and consent to participate

Not applicable.

### Consent for publication

All authors approve the manuscript.

### Availability of data and material

Scripts for generating data plots are available at https://github.com/af-lab/hydractinia. Materials including bacterial vector for Hydractinia histones can be obtain from the laboratory of UF upon request.

### Authors’ contributions

Conceptualization: SGG, UF and AF; Methodology: AT, MGB, TQD, SGG; Investigation: AT, MGB, JCV, IP, TQD, EA, F, SGG; Software: AF; Data Curating: AF, SGG; Writing: AT, SGG, AF, UF; Funding Acquisition, SGG, UF; Supervision, SGG, AF, UF.

## Acknowledgements

We thank members of our labs for lively discussions. Confocal images were taken at the Centre for Microscopy and Imaging Core Facility at NUI Galway.

